# The Drosophila MOZ homolog Enok controls Notch-dependent induction of the RUNX gene *lozenge* independently of its histone-acetyl transferase activity

**DOI:** 10.1101/2020.07.27.222620

**Authors:** Thomas Genais, Delhia Gigan, Benoit Augé, Douaa Moussalem, Lucas Waltzer, Marc Haenlin, Vanessa Gobert

## Abstract

The human KAT6 lysine acetyltransferase MOZ has been shown to be an essential player in the field of normal and malignant hematopoiesis. It belongs to a highly conserved family of epigenetic factors and remodels chromatin by acetylating histone tails in association with its partners of the ING5 complex. Here, we report that its Drosophila counterpart Enok is required during larval hematopoiesis to control the Notch-dependent induction of circulating crystal cells. In particular *enok* is essential to allow expression of the RUNX factor Lozenge (Lz) that controls the crystal cell specific transcriptional program. We demonstrate that this function involves neither the Eaf6 and Ing5 subunits of the Drosophila ING5 complex, nor Enok own acetyltransferase activity. We identify in *lz* third intron a hematopoietic enhancer, which is both required to promote expression in Notch-activated crystal cell precursors in an *enok*-dependent manner and bound by Enok. The non-catalytic mode of action of Enok is likely conserved in MOZ/KAT6 proteins and might be of high relevance in mammalian hematopoiesis, whether normal or malignant.

## Introduction

Hematopoiesis, the highly dynamic process of all blood cell production, is tightly regulated by key transcription and epigenetic factors (Menegatti *et al.* 2019; Fujiwara, 2017; Hu & Shilatifard, 2016). Many of these regulators have been identified through the pathologies induced by their deregulation/mutation. In particular, genetic aberrations such as chromosomal rearrangements have been instrumental, since they allowed immediate identification of the targeted loci. As an example, characterization of the leukemogenic translocation t(8;21) led to the discovery of the transcription factor RUNX1 that was further shown to be essential for multiple steps of mammalian hematopoiesis (Lam & Zhang, 2012). Similarly, the gene encoding Monocytic Leukemia Zinc finger protein (MOZ, also known as KAT6A) was originally identified as the target of myeloid leukemia associated rearrangements, which fuse it to Lysine Acetyltransferases (KAT) such as CREB-binding protein (CBP) or p300, or to the coactivator TIF2 (reviewed in (Katsumoto *et al.* 2008)).

MOZ/KAT6A belongs to a family of KATs conserved from yeast to human, the MYST family (named after its founding members MOZ/Ybf2/Sas2/Tip60), and is mainly known for its ability to acetylate histone H3 on lysine-9, lysine-14 and lysine-23 residues (Voss *et al.* 2009; Qiu *et al.* 2012; Dreveny *et al.* 2014; Lv *et al.* 2017). It is therefore proposed to control transcriptional events by participating in chromatin remodeling *via* the modification of epigenetic marks. MOZ exerts its acetylation activity in a tetrameric ING5 complex also containing the bromodomain PHD finger protein 1 (BRPF1), the human Esa1-associated factor 6 homolog (hEAF6), and the inhibitor of growth 5 (ING5) subunits (Doyon *et al.* 2006; Ullah *et al.* 2008). In mouse, the analysis of a MOZ mutant specifically deprived of its catalytic activity (Perez-Campo *et al.* 2009) reveals a drastic reduction in hematopoietic stem cell (HSC) and committed precursor populations, directly resulting from their reduced proliferation capacities. These results highlight the essential role of MOZ-driven acetylation in HSCs. However, the phenotypic differences existing between this catalytically inactive MOZ mutant and a loss of the entire MOZ protein (Katsumoto *et al.* 2006) suggest that MOZ may also ensure non-acetyltransferase functions *in vivo*, consistent with previous observations that MOZ interaction with RUNX1 promotes transcription independently of its catalytic domain in murine cultured cells (Kitabayashi *et al.* 2001).

Enoki mushroom (Enok), *Drosophila melanogaster* MOZ homolog, was first identified for its role in neuroblast proliferation (Scott *et al.* 2001). Its loss of function results in atrophied mushroom bodies in the fly brain, earning the gene its name, as well as a developmental delay and lethality at the early pupal stage. *enok* was later shown to be required for visual system wiring (Berger *et al.* 2008) and for germline stem cell maintenance, a process during which it ensures both cell autonomous and non-autonomous functions (Xin *et al.* 2013). The first insights into the molecular mechanisms underlying the different roles of Enok came from a study that demonstrated its ability to acetylate histone H3 on lysine-23 (H3K23) (Huang *et al.* 2014); in addition, the authors showed that *enok* is required for the expression of oocyte-polarizing genes and proposed that enok-mediated H3K23 acetylation of its targets promotes their transcriptional activation. Finally, a second study by the same group, demonstrated that Drosophila homologs of the mammalian ING5 complex Br140, Eaf6 and Ing5 interact physically and functionally with Enok for H3K23 acetylation, and identified a new role for Enok in cell cycle regulation (Huang *et al.* 2016). Of note, no function for Enok in hematopoiesis has ever been documented.

*Drosophila melanogaster* is a well-established model to study both normal and pathologic hematopoiesis, since its much simpler hematopoietic system displays a high degree of conservation, whether at the functional or at the molecular level (for an exhaustive review, see (Banerjee *et al.* 2019)). Drosophila blood cells (called hemocytes) are formed in two temporally distinct waves. The embryonic wave gives rise to two differentiated cell types: plasmatocytes that mainly ensure phagocytic functions, and crystal cells that are responsible for melanization reactions and therefore contribute to wound healing and immune response. During larval life, a second wave of hematopoiesis in the lymph gland yields a reservoir of plasmatocytes and crystal cells that remain separated from the circulation until they are released at the onset of metamorphosis (under normal conditions) or mobilized in response to a parasitic infestation (Letourneau *et al.* 2016). Cells formed during the embryonic wave persist in circulation throughout larval stages until adulthood (Holz *et al.* 2003). During larval life, a massive expansion of the plasmatocyte population occurs in clusters of cells located beneath the epidermis (Makhijani & Brückner, 2012). Also, recent live-imaging of sessile hemocytes within these clusters shows that, upon Notch signaling pathway activation, some cells start to express the crystal cell lineage-specific RUNX transcription factor Lozenge (Lz), thus ensuring *de novo* larval crystal cell formation (Leitão & Sucena, 2015).

Here we provide the first evidence that like its mammalian counterpart, Enok participates in hematopoiesis, since we show that the development of larval circulating crystal cells is hindered in *enok* mutants. In particular, we demonstrate that the expression of *lz* depends on a cell-autonomous function of Enok in Notch-activated crystal cell precursors. Unexpectedly, we establish that Enok is required independently of its KAT activity and of the ING5 acetylation complex and we provide the prime example of a catalytic-independent function for a MOZ protein *in vivo*. Finally, our results strongly suggest that Enok directly controls *lz* transcription by binding a key enhancer element participating in its expression in crystal cell precursors.

## Results

### *enok* is essential for circulating larval crystal cell formation

While *enok* functions in various developmental processes were already described (Scott *et al.* 2001; Xin *et al.* 2013; Huang *et al.* 2014, 2016), its possible implication in blood cell formation was not reported. Given the major role played by its homolog MOZ during mammalian hematopoiesis, we decided to examine blood cell formation in various *enok* loss of function mutants. The three different alleles tested (*enok*^*1*^, *enok*^*2*^ and *enok*^*Q253*^), placed either in hemizygote conditions (combined to the *Df(2R)BSC155* deficiency that uncovers the *enok* locus) or in allelic combination (*enok*^*1*^/*enok*^*Q253*^), exhibited a dramatic loss of circulating crystal cells at the larval stage. Functional crystal cells of third instar larvae can be easily visualized, as heat exposition triggers melanization within these cells, which then appear as little black dots. We observed almost no melanized crystal cells in the four *enok* mutant contexts tested as compared to control situations (Fig. 1A), indicating a defect in lineage specification or in precursor cell differentiation. We next examined *enok* mutant larvae circulating blood cells using a Notch Response Element reporter construct (*NRE-GFP*), whose expression is detected in crystal cell precursors and persists until their terminal differentiation (Leitão & Sucena, 2015; Miller *et al.* 2017), and an antibody raised against Prophenoloxidase1 (PPO1), a crystal cell terminal differentiation marker. Our results show that Notch-activated hemocytes (NRE-GFP^+^ cells) have lost PPO1 expression in *enok* loss of function contexts (Fig. 1B), confirming the absence of mature crystal cells in the hemolymph of *enok* loss of function larvae.

**Figure 1.**
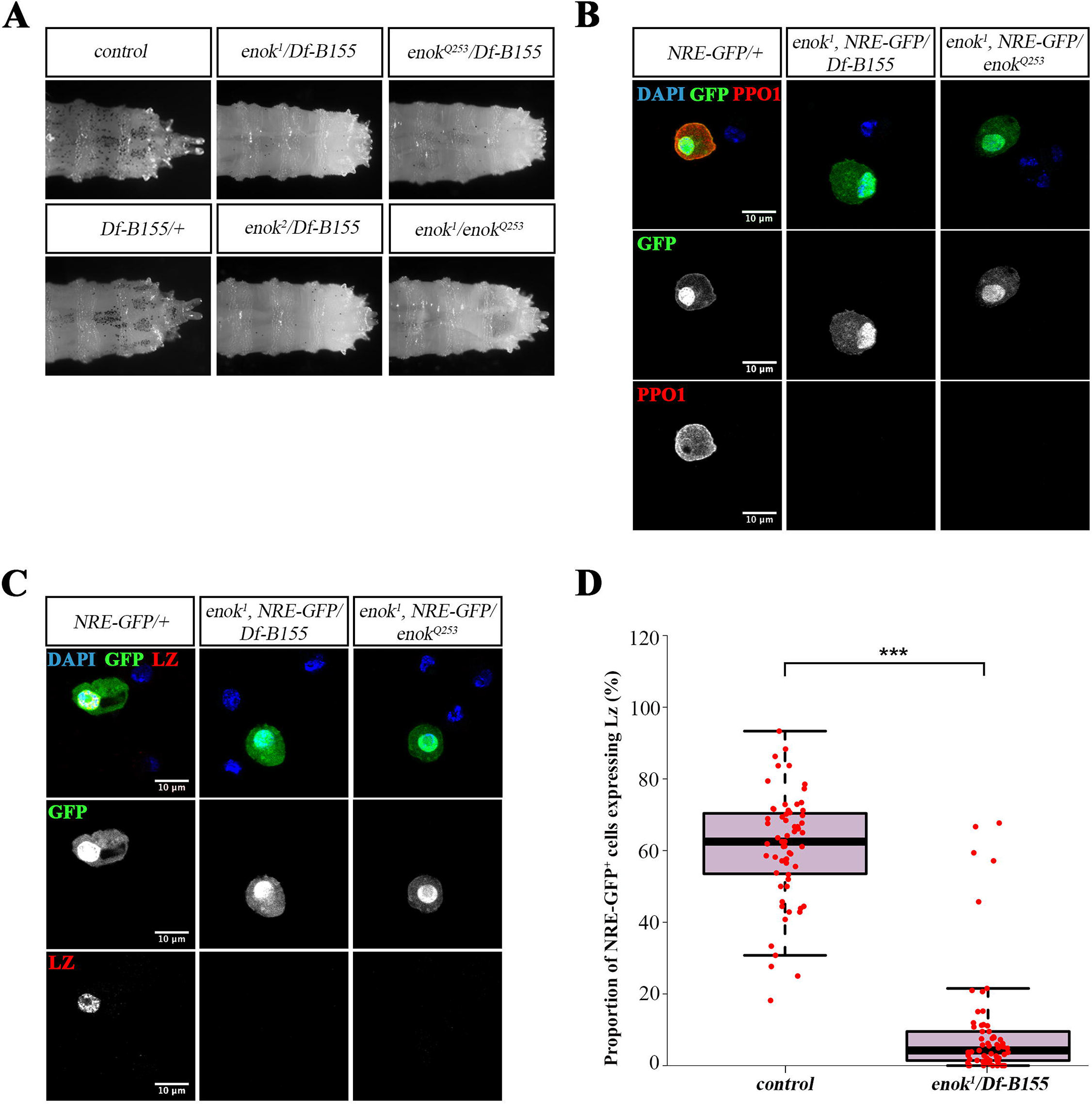
*enok* is required for circulating larval crystal cell formation. (A) Melanization assay on larvae from different *enok* mutant contexts. Posterior side of larvae is to the right. Melanized crystal cells appear as black dots. (B,C) Immunofluorescent staining on circulating cells from larvae of the indicated genotypes (*Df-B155*: abbreviation for *Df(2R)BSC155*); cells were stained with DAPI and either α-PPO1 (B) or α-Lz (C) antibodies. (D) Quantification of the proportions of circulating NRE-GFP^+^ cells expressing Lz as revealed by an immunofluorescent staining. Each dot represents the proportion observed in an individual larva. Complete genotypes: *NRE-GFP*/+ (control) and *enok*^*1*^, *NRE-GFP/Df(2R)BSC155* (enok^1^ /Df-B155). Statistics: **** indicates a p-value≤0.0001.

### *enok* is required for Notch-dependent Lz expression induction

Since the expression of PPO1 depends on the RUNX transcription factor Lz (Waltzer *et al.* 2003; Gajewski *et al.* 2007), we assessed Lz expression in circulating cells and observed that only a small fraction of NRE-GFP^+^ cells retain the ability to express Lz in *enok* loss of function mutants as compared to control larvae (Fig. 1C,D). Interestingly, an increased proportion of Notch-activated hemocytes displays expression of the plasmatocyte specific marker P1 (Supplemental Fig. S1A), which is consistent with the idea that circulating larval crystal cells arise, at least in part, from transdifferentiating plasmatocytes (Leitão & Sucena, 2015). We then evaluated the consequences of *enok* loss of function on other known targets of the Notch signaling pathway (NRE-GFP reporter gene, Su(H)GBE-Gal4,UAS-GFP construct and E(spl)mβ-GFP reporter gene) and did not observe any major deregulation of their expression in *enok* mutant larvae (Supplemental Fig. S1B-D), thus rendering unlikely that the loss of Lz expression results from a generic problem in Notch signaling interpretation. Altogether our results show that *enok* is required downstream of Notch signaling for Lz expression during the formation of circulating larval crystal cells.

### *enok* regulates Lz expression in crystal cell precursors in a cell-autonomous fashion

We next sought to determine in which cells *enok* is required to promote crystal cell production. Directing expression of a *UAS-3HA-enok* transgene in *enok* mutant larvae, either ubiquitously with the *tubulin-Gal4* driver or with the *Notch*^*GMR30A01*^-*Gal4* driver (Miller *et al.* 2017), which allows expression in Notch-activated crystal cell precursors (Supplemental Fig. S2A,B), restores crystal cell maturation (Fig 2A), further demonstrating that the loss of crystal cell linage is due to the loss of *enok* function. In addition, our result indicates that *enok* ensures a cell-autonomous function in the Notch-activated precursors. Furthermore, thorough quantification of the proportion of Lz-expressing cells in circulation confirms that re-expressing *enok* specifically in crystal cell precursors of *enok* mutant larvae is sufficient to restore Lz expression completely (Fig. 2B). We thus conclude that *enok* controls the circulating larval crystal cell lineage formation through a cell-autonomous regulation of Lz expression in Notch-activated precursors.

**Figure 2.**
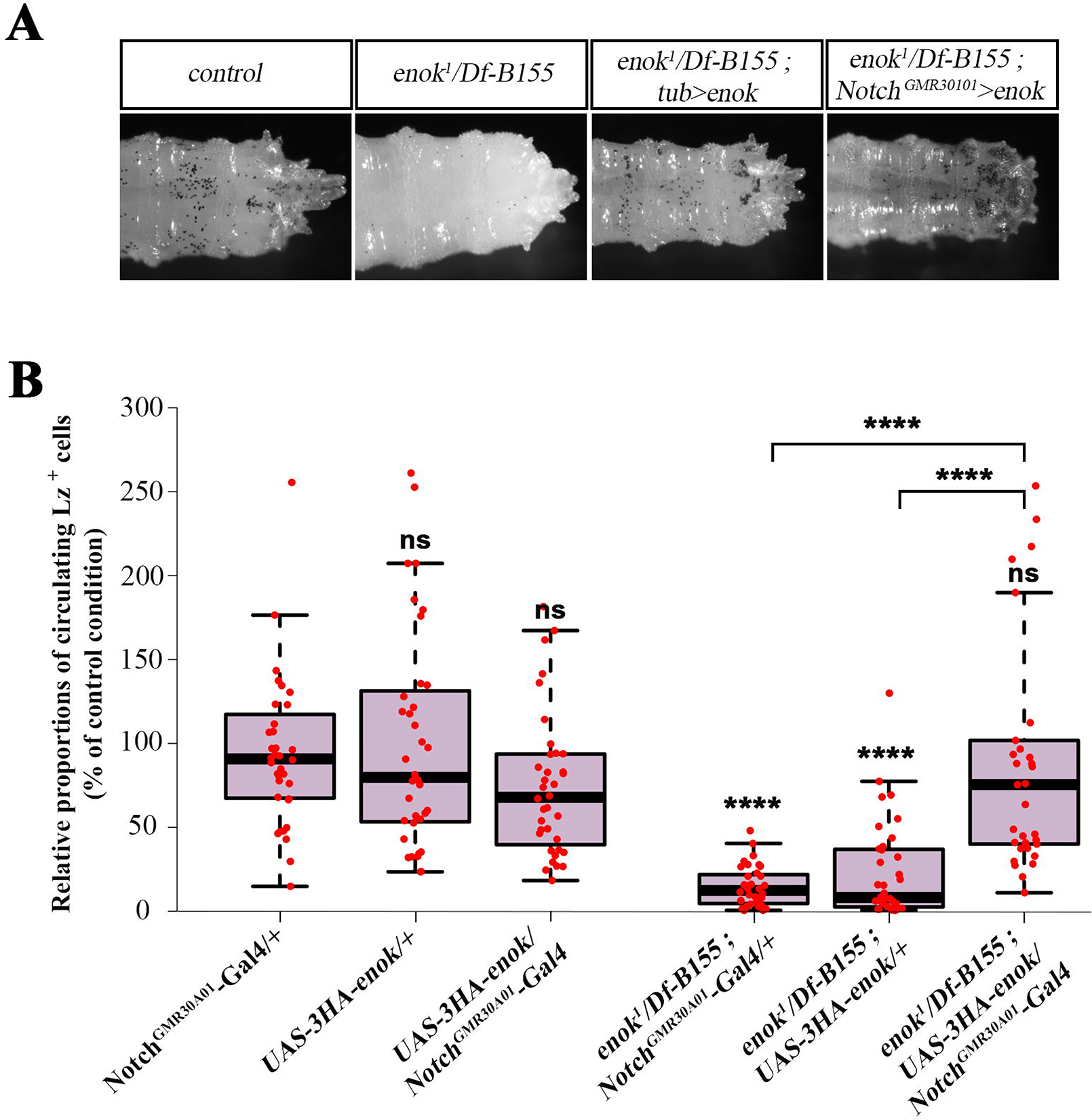
*Notch*^*GMR30A01*^-*Gal4*-driven expression *UAS-3HA-enok* transgene in *enok* mutants restores crystal cell differentiation. (A) Melanization assay on larvae of the indicated genotypes. Complete genotypes: *w*^*1118*^ (control), *enok*^*1*^/*Df(2R)BSC155* (enok^1^/Df-B155), *enok*^*1*^/*Df(2R)BSC155*; *tubulin-Gal4/UAS-3HA-enok* (enok^1^/Df-B155; tub>enok) and *enok*^1^/*Df(2R)BSC155*; *Notch*^*GMR30A01*^-*Gal4/UAS-3HA-enok* (enok^1^/Df-B155; Notch^GMR30A01^>*enok*). (B) Quantification of the relative proportions of Lz positive circulating cells as revealed by immunofluorescent staining (expressed as a percentage of that of the control condition). Each dot represents the proportion of Lz^+^ cells in an individual larva. Statistics: *Notch*^*GMR30A01*^-*Gal4*/+ genotype was used as a reference sample; **** indicates a p-value≤0.0001, ns: not significantly different from reference sample.

### The catalytic activity of Enok and the ING5 complex are dispensable for crystal cell formation

Since Enok is known to cooperate with its partners in the Drosophila ING5 complex to acetylate H3K23 (Huang *et al.* 2014, 2016), we surmised that Enok and the whole ING5 complex might control *lz* expression in Notch-activated hemocytes *via* histone acetylation. While a *Br140*^*S781*^ loss of function allele already existed, no mutation for *Eaf6* or *Ing5* was available. We thus used the CRISPR/Cas9 genome-editing system to excise most of the coding region of these two genes and recovered the *Eaf6*^*M26*^ and *Ing5*^*ex1*^ alleles (see supplemental methods). Also, we introduced in the coding sequence of *enok* a single K807R amino acid substitution that affects a highly conserved Lysine, therefore abrogating its lysine acetyltransferase function (*enok*^*KAT*^ allele, see supplemental methods). As expected, the level of H3K23 acetylation in circulating hemocytes is strongly reduced in *enok*^*1*^ and *Br140*^*S781*^ hemizygote contexts and more mildly in *Eaf6*^*M26*^ hemizygote or *Ing5*^*ex1*^ homozygote mutants (Supplemental Fig. S3). The *enok*^*KAT*^ mutation reduces the level of H3K23 acetylation to that observed with the *enok* null mutant context (*enok*^*1*^/*Df(2R)BSC155)* (Supplemental Fig. S3), suggesting that *enok*^*KAT*^ is a genetically null allele for the acetyltransferase function. It is noteworthy however that, while the levels of H3K23 acetylation are drastically affected in the *enok*^*KAT*^ context, these mutants do not display the developmental delay and lethality that are associated to the other known *enok* mutant loss of function alleles; the *enok*^*KAT*^ mutant adults appear morphologically normal except for a sterility of the females. We then observed in melanization assays that, while *Br140* loss of function results in the same hematopoietic phenotype as *enok* loss of function, gene excision of *Eaf6* or *Ing5* did not affect larval crystal cell formation (Fig. 3A). More strikingly, *enok*^*KAT*^ mutant larvae exhibited seemingly normal crystal cell differentiation (Fig. 3A). Additional quantification of the proportion of NRE-GFP^+^ cells expressing Lz confirms that both *Br140*^*S781*^ and *enok*^*1*^ mutations prevent Notch-activated precursors from differentiating further, while *enok*^*KAT*^ mutation has no effect on this lineage (Fig. 3B). Thus, our results demonstrate that Enok and Br140 regulate Lz through a non-canonic mode of action that is independent of the KAT activity of Enok and of the ING5 complex.

**Figure 3.**
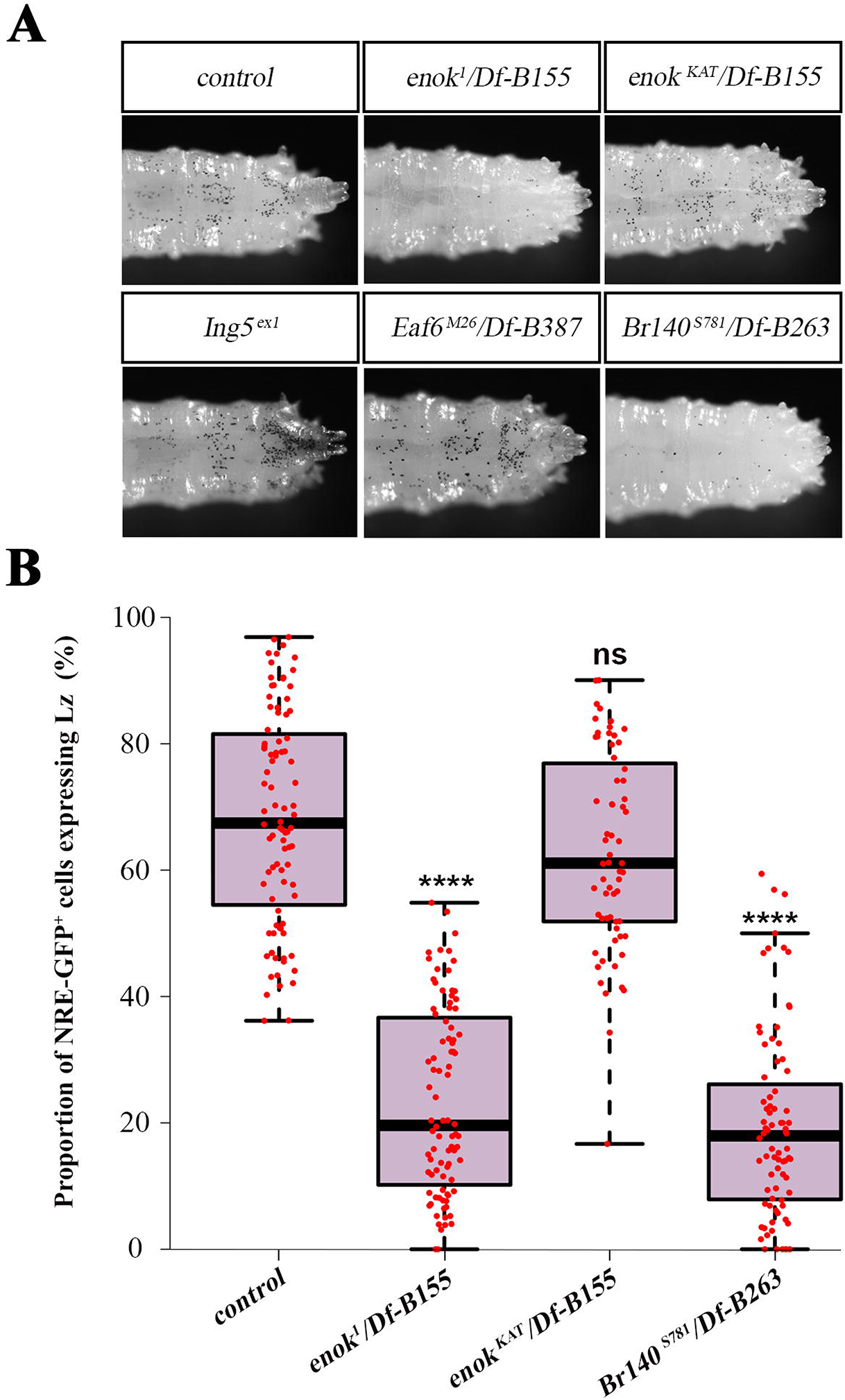
*enok* and *Br140* are required independently of the ING5 complex for crystal cell differentiation. (A) Melanization assay on larvae of the indicated genotypes (*Df-B387*: abbreviation for *Df(3L)BSC387*, *Df-B263*: abbreviation for *Df(2R)BSC263)*. (B) Quantification of the proportions of NRE-GFP^+^ cells expressing Lz as revealed by immunofluorescent staining. Each dot represents the proportion of Lz^+^ cells in an individual larva. Complete genotypes: *NRE-GFP*/+ (control), *enok*^*1*^, *NRE-GFP/Df(2R)BSC155* (enok^1^/Df-B155), *Br140*^*S781*^, *NRE-GFP/Df(2R)BSC263* (Br140^S781^/Df-B263) and *enok*^*KAT*^, *NRE-GFP/Df(2R)BSC155* (enok^KAT^ /Df-B155). Statistics: control genotype was used as a reference sample; **** indicates a p-value≤0.0001, ns: not significantly different from reference sample.

### Lz expression precedes that of Yorkie during circulating larval crystal cell formation

During *de novo* hematopoiesis in the lymph gland, the onset of *lz* expression was reported to be under the control of the Hippo signaling pathway effector, Yorkie (Yki), following Notch signaling activation (Milton *et al.* 2014; Ferguson & Martinez-Agosto, 2014b, 2014a). We reasoned that the same mechanism could be at work in the larval sessile/circulating compartment and that Enok might regulate Yki expression. Our results show that most Notch-activated circulating cells indeed express Yki in a control genetic context and that this expression is lost in *enok* mutant conditions (Fig. 4A,B), indicating that *enok* is also required for Yki expression. Surprisingly however, in control larvae the vast majority of cells expressing both NRE-GFP and Yki also express Lz (Supplemental Fig. S4A), whereas a subset of cells that co-express NRE-GFP and Lz appear negative for Yki staining (Supplemental Fig. S4B, dotted ellipse). The existence of this subpopulation does not fit the model developed for crystal cell differentiation in the lymph gland and raises the possibility that the transcriptional cascade controlling circulating larval crystal cell differentiation might be different. Quantification of Yki expression intensity in *lz*^*R1*^ null mutant larvae shows that crystal cell precursors deprived of *lz* do not express Yki, as compared to a control situation (Fig. 4C), indicating that *lz* is required for Yki expression in these cells. Furthermore, ectopically expressing *lz* in larval circulating hemocytes with the *hemolectin-gal4* driver is sufficient to induce a significant increase of the Yki expressing cell population (Fig. 4D). In contrast, ectopic expression of Yki or activated-Yki (Yki^ACT^) does not trigger Lz expression in circulating cells (Fig. 4D). Altogether our results suggest that following Notch signaling pathway activation in circulating cells, *lz* expression precedes that of *yki*, and is initiated independently of the latter. Therefore, we conclude that the effect of *enok* loss of function on Yki expression is consecutive to the loss of Lz expression in this context. Accordingly, we subsequently focused on the control of Lz expression by *enok*.

**Figure 4.**
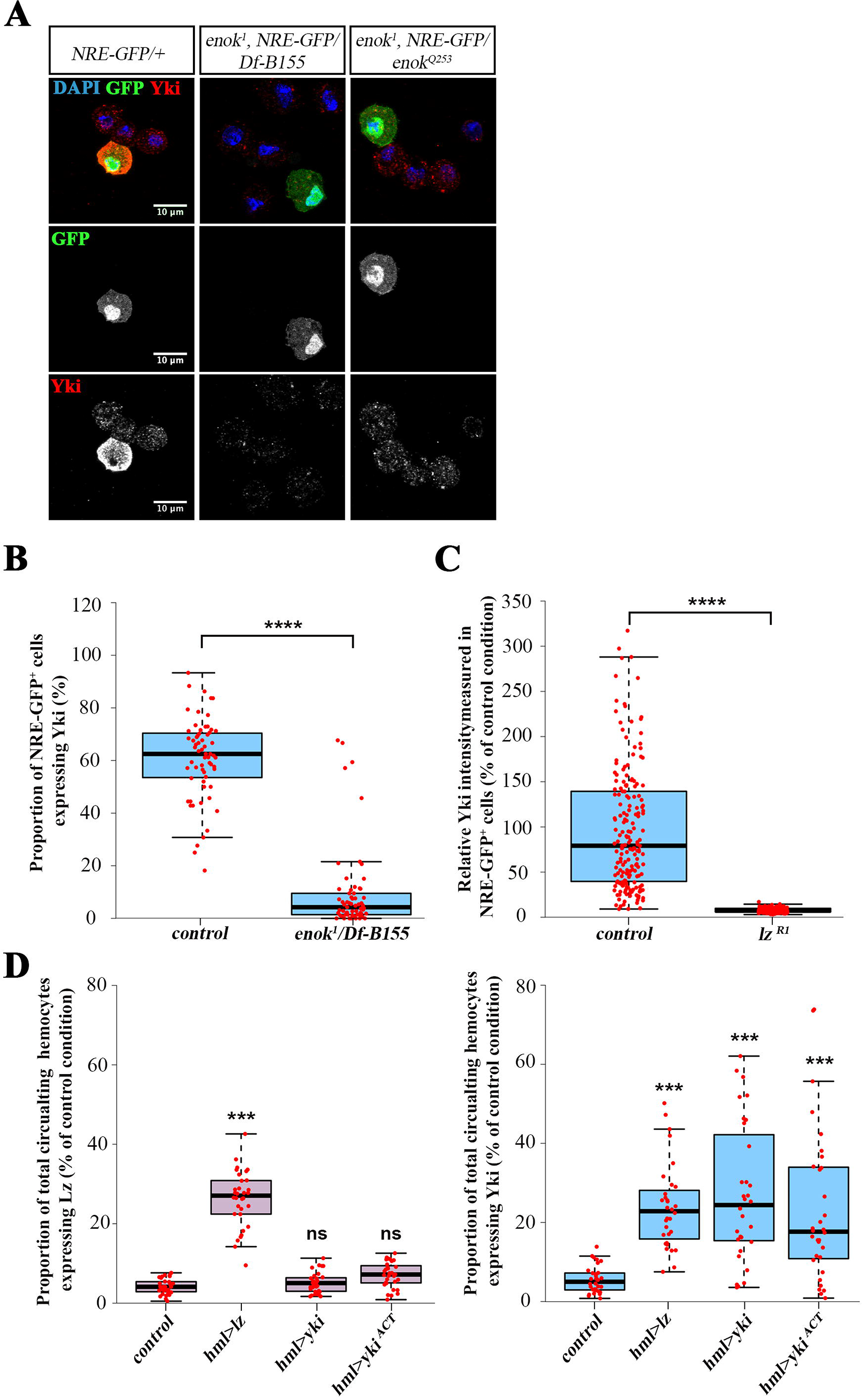
Lz precedes Yki expression during circulating larval crystal cell formation. (A) Immunofluorescent staining on circulating cells from larvae of the indicated genotypes; cells were stained with DAPI and α-Yki. (B) Quantification of the proportions of NRE-GFP^+^ cells expressing Yki as revealed by immunofluorescent staining. Complete genotypes: *NRE-GFP*/+ (control) and *enok*^*1*^, *NRE-GFP/Df(2R)BSC155* (enok^1^/Df-B155). (C) Fluorescence intensity of Yki staining in NRE-GFP^+^ cells (relative to the mean Yki intensity measured in all NRE-GFP^+^ cells of control larvae) in larvae of the following genotypes: *w*^*1118*^/*Y*; *NRE-GFP*/+ (control) and *lz*^*R1*^/*Y*; *NRE-GFP*/+ (lz^R1^). (D) Overexpression of *UAS*-*lz*, *UAS*-*yki* or *UAS*-*yki*^*ACT*^ using the *hml*^*Δ*^-*Gal4* driver. Left panel: proportions of total circulating hemocytes expressing Lz. Right panel: proportions of total circulating hemocytes expressing Yki. Complete genotypes are as follows: *w*/*w*; *hml*^*Δ*^-*Gal4*/+ (control), *w*/*UAS*-*lz*; *hml*^*Δ*^-*Gal4*/+ (UAS-lz), *w*/*w*; *hml*^*Δ*^-*Gal4/UAS-yki-GFP* (UAS-yki) and *w*/*w*; *hml*^*Δ*^-*Gal4*/+; *UAS-yki*^*S168A*^-*GFP*/+ (UAS-yki^ACT^). (B,D) Each dot represents the proportion observed in an individual larva. (C) Each dot represents the intensity value attributed to a single cell. (B-D) Statistics: control genotype was used as a reference sample; **** indicates a p-value≤0.0001, *** indicates a p-value≤0.0010, ns: not significantly different from reference sample.

### Enok binds an intronic region of *lz*, which is required for its expression in circulating crystal cell progenitors

We selected a set of transgenic lines containing potential regulatory regions of *lz* placed upstream of the Gal4 coding sequence and tested them for their ability to recapitulate *lz* expression in the NRE-GFP expressing cells (Supplemental Table 1). Among these, the 2 kb regulatory region covered by the Vienna Tile VT059215 (Fig. 5A), either placed upstream of the Gal4 coding sequence (*lz*^*VT059215*^-*Gal4*, Supplemental Fig. 5A) or directly fused to the RedStinger reporter gene (*lz*^*VT059215*^-*RedStinger*, Supplemental Fig. 5B) is able to drive transcription in a large fraction of the NRE-GFP expressing cells, which overlaps importantly but not exclusively with endogenous Lz expression (Supplemental Fig. 5A,B). This potential regulatory region is embedded in the third intron of the *lz* gene (Fig. 5A). In order to establish the role played by this region, we deleted *lz* third intron (5 kb) using the CRISPR/Cas9 genome editing technique (*lz*^*int3*^ allele (Fig. 5A), see supplemental methods). Despite the critical roles of *lz* during eye development (Daga *et al.* 1996), the excision of *lz* third intron yields individuals with normal eyes, indicating that in these mutants, *lz* expression in the eye imaginal disc is not affected and that the Lz protein is functional. However, like *enok* loss of function, *lz*^*int3*^ mutation abolishes Lz expression in most Notch-activated hemocytes (Fig. 5B,C), indicating that *lz* third intron contributes to its expression in circulating crystal cell precursors. In addition, *lz*^*VT059215*^-*Gal4* driven re-expression of *lz* in a *lz*^*R1*^ mutant background is sufficient to restore a wild-type proportion of Lz expressing cells in the NRE-GFP domain (Fig. 5E), as well as crystal cell differentiation (Fig. 5D). Altogether, these results show that *lz* third intron contains a hematopoietic enhancer both necessary and sufficient for its expression in most crystal cell precursors. We then speculated that Enok might occupy this region to promote Lz expression in Notch-activated hemocytes. Chromatin immunoprecipitation performed in *tub-Gal4/UAS-3HA*-*Enok* embryos revealed a conspicuous enrichment in HA-Enok binding across regions of *lz* third intron, corresponding to the identified VT059215 regulatory element (Fig. 5F), and a modest (yet statistically significant) binding of HA-Enok in *lz* first exon, which was recently reported to be bound by Br140 (Kang *et al.* 2017), suggesting potential co-occupancy of this region by an Enok/Br140 complex. No obvious enrichment was detected in the upstream intergenic region previously reported to contain some *lz* regulatory elements (Bataillé *et al.* 2005; Ferjoux *et al.* 2007). These results indicate that the *lz*^*VT059215*^ region might be an important site of Enok binding in *lz* and prompted us to test whether the expression driven by this enhancer depends on Enok or not. We thus quantified the expression of the *lz*^*VT059215*^-*RedStinger* reporter gene in Notch-activated circulating cells, in the presence or the absence of Enok, and observed that its expression level is indeed reduced in two different *enok* loss of function conditions as compared to control (Fig. 5G). Altogether, our results show that the third intron of *lz* contains a regulatory region that is essential for *its* expression in Notch-activated hemocytes and that is regulated by Enok.

**Figure 5.**
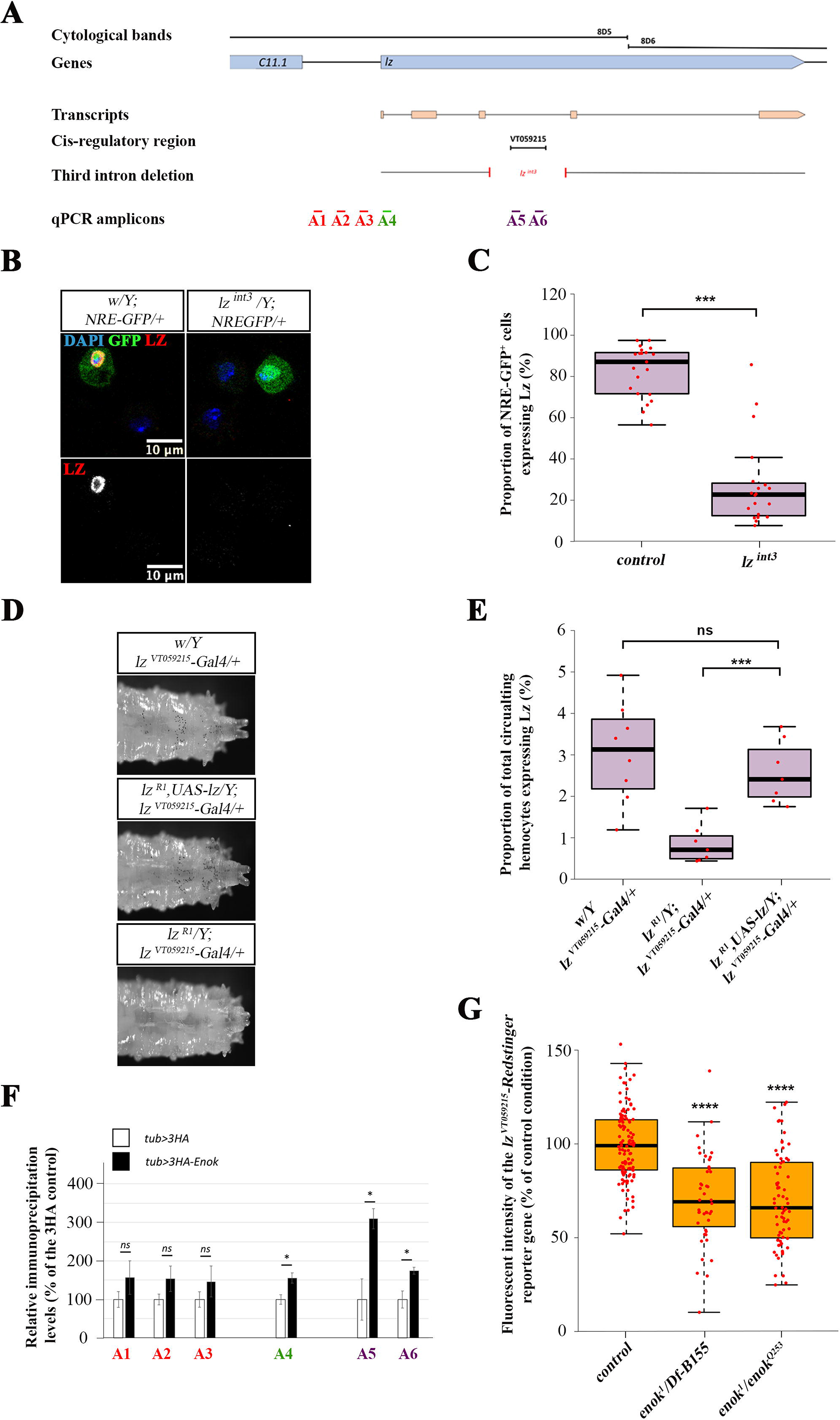
Enok controls a hematopoietic enhancer in *lz* third intron. (A) Schematic representation the *lz* locus and the different genetic elements or reagents used in this study. (B) Immunofluorescent staining on circulating cells from *NRE-GFP*/+; *lz*^*VT059215*^-*Gal4*,*UAS*-*RedStinger*/+ larvae; cells were stained with DAPI and α-Lz antibody. (C) Quantification of the proportion of NRE-GFP^+^ cells expressing Lz in *lz*^*int3*^ and control male larvae, as revealed by immunofluorescent staining. Each dot represents the proportion of Lz^+^ cells in an individual larva. (D) Melanization assay on larvae of the indicated genotypes. (E) Quantification of the proportion of total hemocytes expressing Lz as revealed by immunofluorescent staining. Each dot represents the proportion of Lz^+^ cells in an individual larva. (F) Chromatin Immunoprecipitation and qPCR performed on *tub-Gal4/UAS-3HA* (3HA, white) and *tub-Gal4/UAS-3HA*-*enok* whole embryos (3HA-*Enok*, black) using a α-HA antibody. (G) Quantification of the fluorescence intensity of the *lz*^*VT059215*^-Redstinger reporter gene in control and *enok* mutant conditions. Each dot represents the mean fluorescence intensity measured in cells from a single larva. Statistics: (C,G) control genotype was used as a reference sample, (E) *w*/*Y*; *lz*^*VT059215*^-*Gal4*/+ genotype was used as a reference sample, (C,E,F,G) **** indicates a p-value≤0.0001, *** indicates a p-value≤0.0010, * indicates a p-value≤0.0500, ns: not significantly different from reference sample.

## Discussion

In the present study, we characterize the effect of *enok* loss of function on the circulating crystal cell lineage and demonstrate that *enok* is required in a cell-autonomous manner in Notch-activated hemocytes during the formation of circulating larval crystal cells. In particular, we present evidence that Enok acts downstream of the Notch signaling pathway to ensure expression of the crystal cell fate determinant Lz in crystal cell precursors and that it relies on a KAT-independent mechanism to do so. Finally, our findings pinpoint a *cis*-regulatory region critical for *lz* expression in circulating crystal cell precursors, which is both bound and regulated by Enok.

Our most striking result is that the control of Lz expression relies on a catalytic-independent mode of action of Enok likely involving its interactor Br140, but for which Eaf6 and Ing5 appear to be dispensable. The *enok* mutant alleles used in previous studies cause a developmental delay and a general lethality since 50% of mutant animals die before puparium formation and those that pupate never emerge as adult (Scott *et al.* 2001). These phenotypes are not developed by catalytically-dead mutants, indicating that H3K23 acetylation is largely dispensable in these processes. The observed sterility of *enok*^*KAT*^ mutant adult females is consistent with the idea that H3K23 acetylation by Enok is essential for the oocyte polarization (Huang *et al.* 2014) and suggests that its function in the maintenance of germline stem cells (Xin *et al.* 2013) requires its catalytic activity. Catalytic-independent functions of MOZ/KAT6A, the mammalian homolog of Enok, have been speculated based on the phenotypic differences observed between mice either totally deprived of MOZ (Katsumoto *et al.* 2006), or in which its catalytic activity has been abolished by a single amino acid substitution (Perez-Campo *et al.* 2009). However, there is so far no documented study of such KAT-independent functions of MOZ *in vivo*. Our work provides the first defined example of a catalytic-independent contribution of a MOZ/KAT6 protein in an integrated model, therefore bringing the proof of principle that these proteins rely on a dual mode of action (catalytic and non-catalytic) to ensure their various functions. Yet this new operating mode remains to be elucidated at the mechanistic level.

In mammals MOZ has been shown to interact with a variety of transcription factors (such as RUNX1, PU.1, c-JUN, P53, TEL, NF-κB, NRF2, reviewed in (Perez-Campo *et al.* 2013)) to regulate target gene expression and in particular, its role as a transcriptional coactivator of RUNX1 in cell culture (Kitabayashi *et al.* 2001; Bristow & Shore, 2003) was shown not to depend on its catalytic domain (Kitabayashi *et al.* 2001). Thus, like MOZ, Enok could interact with a yet to be identified transcription factor on the VT059215 enhancer to control *lz* expression in Notch-activated circulating hemocytes. The VT059215 regulatory region contains elements that are relevant to the regulation of *lz*, including binding sites for Su(H) (the effector of Notch signaling pathway), for Scalloped (the major partner of Yki), for GATA factors (such as Serpent, a major regulator of blood cells development) and for Lz itself. Since Lz is the homolog of mammalian RUNX1 and is already known to participate in an autoregulatory loop during embryogenesis (Ferjoux *et al.* 2007), an Enok/Lz interaction on the VT059215 enhancer could participate in a feed-forward loop allowing *lz* expression maintenance in circulating larval crystal cells. However, neither the MOZ-interacting domain of RUNX1 nor the C-terminal RUNX1-interacting domain of MOZ are conserved in their respective Drosophila counterparts, rendering this hypothesis less likely. Further characterization of the requirement for the different binding sites in the VT059215 regulatory region could allow the identification of the transcription factors regulating *lz* expression *via* this enhancer and whose properties would be regulated by Enok.

Umer *et al.* demonstrated in a recent study that Enok behaves as a Trithorax group (TrxG) protein; they showed that *enok* loss-of-function suppresses the phenotype induced by the downregulation of Polycomb (Pc), a major epigenetic silencer, and that Enok antagonizes Pc recruitment on chromatin at TrxG target gene loci (Umer *et al.* 2019). Work by Strübbe *et al.* showed that in Drosophila embryos Enok and Br140 interact physically with Pc (Strübbe *et al.* 2011), which was further confirmed in another study highlighting the substantial overlap between chromatin occupancies by Br140 and by the Polycomb Repressive Complex 1, PRC1 (Kang *et al.* 2017) on developmentally regulated bivalent genes. Of note, the authors did not observe any enrichment in the Enok-dependent H3K23ac mark on their set of bivalent genes, which is compatible with a non-catalytic mode of action of Enok. Interestingly, one of the genomic regions co-occupied by Br140 and PRC1 is the *lz* locus. Binding of the Enok/Br140 pair on the VT059215 regulatory region could therefore contribute to limit the excessive silencing of the gene, thus allowing its expression in response to developmental cues such as Notch signaling pathway activation. In the absence of Enok or Br140, the locus would subsequently become mostly unresponsive to Notch signaling.

In conclusion, we have shown that like MOZ in mammals, Enok is an important regulator of hematopoiesis in Drosophila and we demonstrated that its function in physiological hematopoiesis relies on a non-catalytic activity. It seems very important to fully elucidate this KAT-independent mode of action of MOZ/KAT6 proteins, as it might be of high relevance in the development of MOZ-related pathologies as well as during normal hematopoiesis.

## Material and methods

### Drosophila stocks, genetics and melanization assays

Stocks and crosses were maintained at 25°C on standard yeast-agar-cornmeal medium. The complete list of stocks used in this study is provided as supplemental material (Supplemental Table 2). Larvae collected for phenotypic analysis were raised in controlled density conditions: in brief, for each cross, 10 females were fertilized by 4 males and left to lay eggs for 16 hours at 25°C. Third instar wandering larvae of the appropriate genotypes were collected 5 days after egg laying. Female larvae were used, unless otherwise specified. For melanization assays, two batches of six larvae were collected in two 1.5 mL eppendorf tubes containing 200 uL of Phosphate Buffered Saline (PBS) each, incubated in a 65°C waterbath for 30 minutes, stored on ice for 30 minutes and imaged on a SMZ18 stereomicroscope (Nikon). Each experiment was reproduced at least three times (minimum of total larvae observed for each phenotype, n=36).

### Immunofluorescent staining and Operetta quantifications on circulating cells

Four female third instar larvae were bled in 1 ml of PBS in 24-well-plate containing a glass coverslip. Hemocytes were centrifuged for 2 min at 900 g, fixed for 20 minutes with 4% paraformaldehyde in PBS and washed twice in PBS. Cells were then permeabilized in PBS-0.3% Triton (PBST), blocked in PBST-1% Bovine Serum Albumin (BSA) and incubated with primary antibodies at 4°C over night in PBST-BSA. The complete list of primary antibodies used in this study is provided as supplemental material (Supplemental Table 3). Next, cells were washed in PBST, incubated for 2 hours at room temperature with corresponding Alexa Fluor-labeled secondary antibodies (Molecular Probes), washed in PBST and mounted in Vectashield medium (Eurobio-Vector) following incubation with DAPI. Imaging was performed on a Leica SP8 confocal microscope. For fluorescence quantification, single female larvae were bled individually in 96-well-plate and samples were processed as described above; imaging was performed on an Operetta microscope (Perkin-Elmer); for each well 30 fields were captured at 20X magnification. Statistical analyses were performed with GraphPad Prism (GraphPad Software, Inc.); two-tailed Student t-Test were used for comparisons of two samples and one-way ANOVA followed by Dunnett test for comparisons of more than two samples.

### Generation of UAS-3HA-Enok and lz^VT059215^-Red-Stinger fly strains

- cloning of UAS-3HA-Enok: a three HA-tags encoding sequence was directionally inserted in the pUAStAttB vector (Addgene) polylinker using EcoRI and NotI restriction enzymes, yielding a pUAStAttb-3HA plasmid. A p-BlueScript containing the full-length *enok* cDNA (kind gift from Takashi Suzuki) was then used as a template to PCR-amplify the *enok* coding sequence (Phusion high-fidelity polymerase, Thermofisher scientific). The amplified fragment was cloned in frame with the 3HA sequence, between the NotI and KpnI restriction sites of the pUAStAttB-3HA vector, using the In-fusion HD cloning kit (Clontech) according to supplier’s instructions.

- cloning of lz^VT059215^-Red-Stinger: this cloning was performed in two steps. We first generated a phiC31-based transformation vector containing a red fluorescent reporter gene compatible with golden gate cloning. Briefly, the pAttB vector (Bischof *et al.* 2013)(kind gift of K. Basler) was modified to remove the existing BsaI restriction sites and we then inserted a polylinker containing BsaI restriction sites on each side of a *lacZ* gene from the pCambia2200 vector (kind gift of Jean-Philippe Combier), the hsp70 promoter and the DsRedT4-NLS coding sequence from the pRed-H-Stinger vector (Barolo *et al.* 2004)(DGRC). In a second step, we cloned the VT059215 sequence into the pAttB-Red-H-Stinger vector with BsaI restriction enzyme following a golden gate cloning protocol.

All constructs were checked by sequencing for polymorphisms/mutations, prior to injection for phiC31-mediated insertion in an AttP2 platform containing fly line. All plasmid sequences and detailed cloning procedures are available on request.

### Chromatin Immunoprecipitation on embryos

Four independent chromatin immunoprecipitation experiments were performed as follows: embryos were collected twice a day for 2-3 days, washed, frozen dry and kept at −80°C. They were then processed as described in (Loubiere *et al.* 2017). The immunoprecipitated chromatin was subsequently used in RealTimePCR using the SsoFast Evagreen chemistry (Biorad) in a CFX96 thermocycler (Biorad). 100 mg of embryos were used for each replicate. The complete list of primers used in this study is provided as supplemental material (SuppTable 4).

## Supporting information

Supplemental figure legends methods and references

## Acknowledgments

We are grateful to Leila Abraham, Camille Calvel, and Maxime Fayel for their participation in mutant fly strain engineering, Amélie Destenabes and Julien Favier at the CBD fly facilities for transgenesis and CRISPR/Cas9 injections, Michèle Crozatier, Nathalie Vanzo and Caroline Monod for critical reading of the manuscript and all our colleagues at the CBD for appreciated comments at various steps of the project. We thank Brice Ronsin, Stéphanie Bosch and Jessie Bourdeaux at the Toulouse RIO imaging platform for assistance with confocal microscopy and Operetta imaging and Benjamin Tritschler for invaluable help with logistics. We are also deeply grateful to Drs Erjun Ling, Takashi Suzuki, Matthew Gibson, Steven Hou, Konrad Basler and Jean-Philippe Combier for sharing resources, and to the Bloomington stock center, the Vienna Drosophila Ressource Center, the Kyoto stock center and the Developmental Studies Hybridoma Bank for providing us with antibodies and fly stocks. TG was supported by a doctoral fellowship from La Ligue Nationale contre le Cancer, and the project benefitted from the support of the Centre National de Recherche Scientifique (CNRS) and the Université Toulouse III - Paul Sabatier (UPS).

