## Supplemental figure legends methods and references for "The Drosophila MOZ homolog Enok controls Notch-dependent induction of the RUNX gene *lozenge* independently of its histone-acetyl transferase activity"

**Supplemental Figure S1.** Circulating crystal cell precursors display increased plasmatocyte marker expression despite normal Notch signaling pathway activation in an *enok* mutant context. (A) Quantification of the proportions of NRE-GFP+ cells expressing the plasmatocyte specific marker P1 as revealed by an immunofluorescent staining. Statistics: **** indicate a p-value≤0.0001. (B) Quantification of the proportions of total circulating hemocytes expressing the NRE-GFP reporter gene in larvae of the following genotypes: *NRE-GFP/+* (control) and *enok^1^, NRE-GFP/Df(2R)BSC155 (*enok^1^/Df-B155). (C) Quantification of the proportion of total circulating hemocytes expressing the *UAS-GFP* construct under the control of the *Su(H)-GBE-Gal4* driver. Complete genotypes: *UAS-GFP*/+; *Su(H)-GBE-Gal4/+* (control) and *UAS-GFP*/+; *enok^1^/Su(H)-GBE-Gal4*, *Df(2R)BSC155* (enok^1^/Df-B155). (D) Immunofluorescent staining on circulating cells from larvae of the indicated genotypes; cells were stained with DAPI and α-Lz antibody. Complete genotypes: E(spl)*mβ-GFP/+* (control) and E(spl)*mβ-GFP,enok^1^/Df(2R)BSC155* (enok^1^/Df-B155). (B,C) Each dot represents the proportion measured in an individual larva.

**Supplemental Figure S2.** *Notch^GMR30A01^-Gal4* directs expression in the crystal cell lineage. (A) Left: Proportion of Notch^GMR30A01^-Gal4 positive cells expressing the NRE-GFP reporter gene in *NRE-GFP/+; Notch^GMR30A01^-Gal4, UAS-RedStinger larvae*. Middle: Proportion of NRE-GFP positive cells expressing the Notch^GMR30A01^-Gal4 construct in *NRE-GFP/+; Notch^GMR30A01^-Gal4, UAS-RedStinger larvae*. Right: Venn diagram summarizing the repartition of the different cell categories in circulating hemocytes. Each dot represents the proportion observed in an individual larva. (B) Lineage-tracing of the *Notch^GMR30A01^-Gal4* driver. Circulating cells were stained with DAPI and α-Lz antibody. Proportions of each cell categories were quantified in a *UAS-RedStinger, UAS-FLP, Ubi-p63E(FRT.STOP)Stinger/Notch^GMR30A01^-Gal4* genetic context (G-trace system ).

**Supplemental Figure S3.** H3K23 acetylation in circulating hemocytes depends on the ING5 complex and on the catalytic activity of Enok. Mean fluorescence intensity in larval circulating hemocytes (relative to the mean intensity measured in total hemocytes of control larvae), as revealed by an α-acetylated-H3K23 immunostaining. Complete genotypes: *w^1118^/+* (control)*,* *enok^1^/Df(2R)BSC155* (enok^1^/Df-B155), *Br140^S781^/Df(2R)BSC263* (Br140^S781^/Df-B263), *Eaf6^M26^/Df(3L)BSC387* (Eaf6^M26^/Df-B387), *Ing5^ex1^/Ing5^ex1^* (Ing5^ex1^) and *enok^KAT^/Df(2R)BSC155* (enok^KAT^/Df-B155). Each dot represents the value attributed to a single cell. Statistics: control genotype was used as a reference sample; *** indicate a p-value≤0.0010.

**Supplemental Figure S4.** In the NRE-GFP^+^ domain, all Yki-expressing cells also express Lz, whereas some Lz-expressing cells do not express Yki. (A) Relative fluorescence intensity of the NRE-GFP transgene plotted against the relative fluorescence intensity of Yki staining in total hemocytes of control larvae. The horizontal dashed line represents the threshold for NRE-GFP positive cells (upper part) and the vertical dashed line represents the threshold for Yki positive cells (right part). The green-boxed area corresponds to the NRE-GFP/Yki double positive population; blue dots are cells negative for Lz expression and red dots are cells positive for Lz expression. (B) Relative fluorescence intensity of the NRE-GFP transgene plotted against the relative fluorescence intensity of Lz staining in total hemocytes of control larvae. The horizontal dashed line represents the threshold for NRE-GFP positive cells (upper part) and the vertical dashed line represents the threshold for Yki positive cells (right part). The pink-boxed area corresponds to the NRE-GFP/Lz double positive population; blue dots are cells negative for Yki expression and red dots are cells positive for Yki expression. The dashed ellipse encloses NRE-GFP/Lz double positive cells that do not express Yki.

**Supplemental Figure S5.** The *lz^VT059215^* regulatory region promotes expression in a large fraction of Lz^+^ Notch-activated hemocytes. (A) Immunofluorescent staining on circulating cells from *NRE-GFP/+; lz^VT059215^-Gal4,UAS-RedStinger/+* larvae; cells were stained with DAPI and α-Lz antibody. (B) Quantification of the proportions of the different categories of NRE-GFP^+^ cells in *NRE-GFP/+; lz^VT059215-^RedStinger/+* larvae. Each dot represents the proportion of the indicated cell type observed in an individual larva.

**Supplemental methods**

**Generation of mutant fly lines by CRISPR/Cas9-mediated genome editing**

Single guide RNA (sgRNA)-compatible sequences were identified in the regions of interest with the Geneious software (Biomatters Ltd); selected sequences had null off-target scores. Oligonucleotides for sgRNA cloning in the pCFD3 expression vector (Port *et al*, 2014) were synthetized by Integrated DNA Technologies and processed according to: <http://www.crisprflydesign.org/wp-content/uploads/2014/05/Cloning-with-pCFD3.pdf>. Constructs were checked by sequencing prior to injection in *vasa-Cas9* transgenic fly strains (Port *et al*, 2014). Three different editing strategies were used depending on the type of mutant to generate.

*-* Eaf6 *and* Ing5 *genes excisions* *(Non Homologous Ends Joining)*: we took advantage of the presence of P-elements inserted in the vicinity of the 5’UTR of the genes and used a *mini-white* marker reversion-based screening strategy. Briefly, the d06605 transposon (*Eaf6*) or the EY13664 transposon (*Ing5*) was brought in a *vasa-Cas9* genetic background using standard Drosophila genetics. The resulting fly strains were used for injection of two sgRNA-containing pCFD3 vectors targeting each side of the inserted P-element. sgRNA expression vectors were injected at a concentration of 250 ng/uL each. Injected F_0_ individuals were crossed, then their F_1_ progeny was screened for the reversion of the *mini-white* marker and positive F_1_ individuals were used to establish stocks. Gene excisions were confirmed by PCR on genomic DNA extracted from F_2_ flies.

*-* enok^KAT^ *allele engineering (single aminoacid substitution, Homology Directed Repair)*: the K807R mutation introduced in the MYST domain of *enok* affects a Lysine that is highly conserved across evolution and critical for the KAT activity of yeast and human MYST proteins (Yuan *et al*, 2012; Yang *et al*, 2012). A repair DNA template called single stranded oligonucleotide donor (ssODN) was injected along with the sgRNA expression vector so as to direct the reparation (ssODN: 100 ng/uL and sgRNA: 250 ng/uL). The ssODN sequence contained the mutation of interest, as well as a sabotage mutation of the Protospacer Adjacent Motif to avoid re-cleaving of an edited chromosome. In addition, the nucleotide sequence immediately surrounding the desired site of modification was degenerated by introduction of silent mutations, in order to allow the design of a discriminant screening primer. 10-15 F_1_ males were crossed individually for each F_0_ founder; after 4-5 days in the crossing vials, each F_1_ male was retrieved for single fly DNA extraction. The PCR-based screen of F_1_ individuals relied on a triple primer PCR: two external primers amplify DNA in both positive and negative editing events, while an internal primer specifically amplifies DNA if the desired mutation is present.

*-* lz *third intron excision (Non Homologous Ends Joining)*: two sgRNA expression vectors targeting each extremity of *lz* third intron were injected at a concentration of 250 ng/uL each; 10-15 F_1_ males were crossed individually for each F_0_ founder and after 4-5 days in the crossing vials, they were retrieved and tested for their ability to generate shorter amplicons in PCR reactions.

All recovered events were further confirmed and characterized by sequencing of the edited loci. Sequences of sgRNA, ssODN and primers used in the various strategies are provided in SuppTable 4. Mutant characterization sequences are longer and therefore are only available on request.

**Additional References (related to Supplemental Tables and supplemental methods)**

Bellen HJ, Levis RW, Liao G, He Y, Carlson JW, Tsang G, Evans-Holm M, Hiesinger PR, Schulze KL, Rubin GM, Hoskins RA & Spradling AC (2004) The BDGP gene disruption project: single transposon insertions associated with 40% of Drosophila genes. *Genetics* **167:** 761–781

Evans CJ, Olson JM, Ngo KT, Kim E, Lee NE, Kuoy E, Patananan AN, Sitz D, Tran P, Do M-T, Yackle K, Cespedes A, Hartenstein V, Call GB & Banerjee U (2009) G-TRACE: rapid Gal4-based cell lineage analysis in Drosophila. *Nat. Methods* **6:** 603–605

Hazelrigg T, Levis R & Rubin GM (1984) Transformation of white locus DNA in drosophila: dosage compensation, zeste interaction, and position effects. *Cell* **36:** 469–481

Lee T & Luo L (1999) Mosaic Analysis with a Repressible Cell Marker for Studies of Gene Function in Neuronal Morphogenesis. *Neuron* **22:** 451–461

Oh H & Irvine KD (2008) In vivo regulation of Yorkie phosphorylation and localization. *Development* **135:** 1081–1088

Port F, Chen H-M, Lee T & Bullock SL (2014) Optimized CRISPR/Cas tools for efficient germline and somatic genome engineering in Drosophila. *PNAS* **111:** E2967–E2976

Thibault ST, Singer MA, Miyazaki WY, Milash B, Dompe NA, Singh CM, Buchholz R, Demsky M, Fawcett R, Francis-Lang HL, Ryner L, Cheung LM, Chong A, Erickson C, Fisher WW, Greer K, Hartouni SR, Howie E, Jakkula L, Joo D, et al (2004) A complementary transposon tool kit for Drosophila melanogaster using P and piggyBac. *Nat Genet* **36:** 283–287

Tracey WD, Ning X, Klingler M, Kramer SG & Gergen JP (2000) Quantitative analysis of gene function in the Drosophila embryo. *Genetics* **154:** 273–284

Yang C, Wu J, Sinha SH, Neveu JM & Zheng YG (2012) Autoacetylation of the MYST lysine acetyltransferase MOF protein. *J. Biol. Chem.* **287:** 34917–34926

Yuan H, Rossetto D, Mellert H, Dang W, Srinivasan M, Johnson J, Hodawadekar S, Ding EC, Speicher K, Abshiru N, Perry R, Wu J, Yang C, Zheng YG, Speicher DW, Thibault P, Verreault A, Johnson FB, Berger SL, Sternglanz R, et al (2012) MYST protein acetyltransferase activity requires active site lysine autoacetylation. *EMBO J.* **31:** 58–70

Zeng X, Chauhan C & Hou SX (2010) Characterization of midgut stem cell- and enteroblast-specific Gal4 lines in drosophila. *Genesis* **48:** 607–611
