## Supplementary figures and images for "The Drosophila MOZ homolog Enok controls Notch-dependent induction of the RUNX gene *lozenge* independently of its histone-acetyl transferase activity"

### Supplemental Figure 1

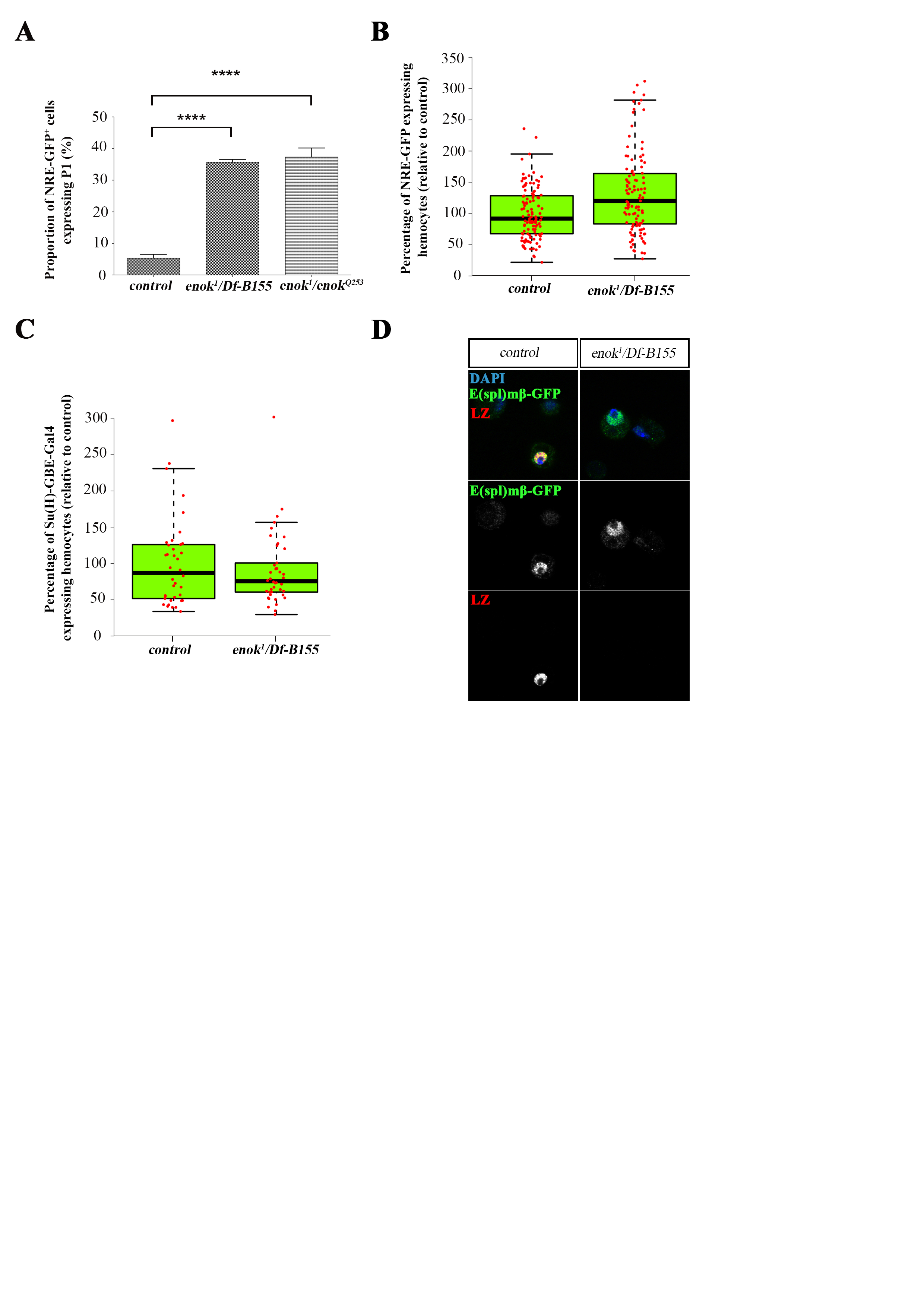

### Supplemental Figure 2

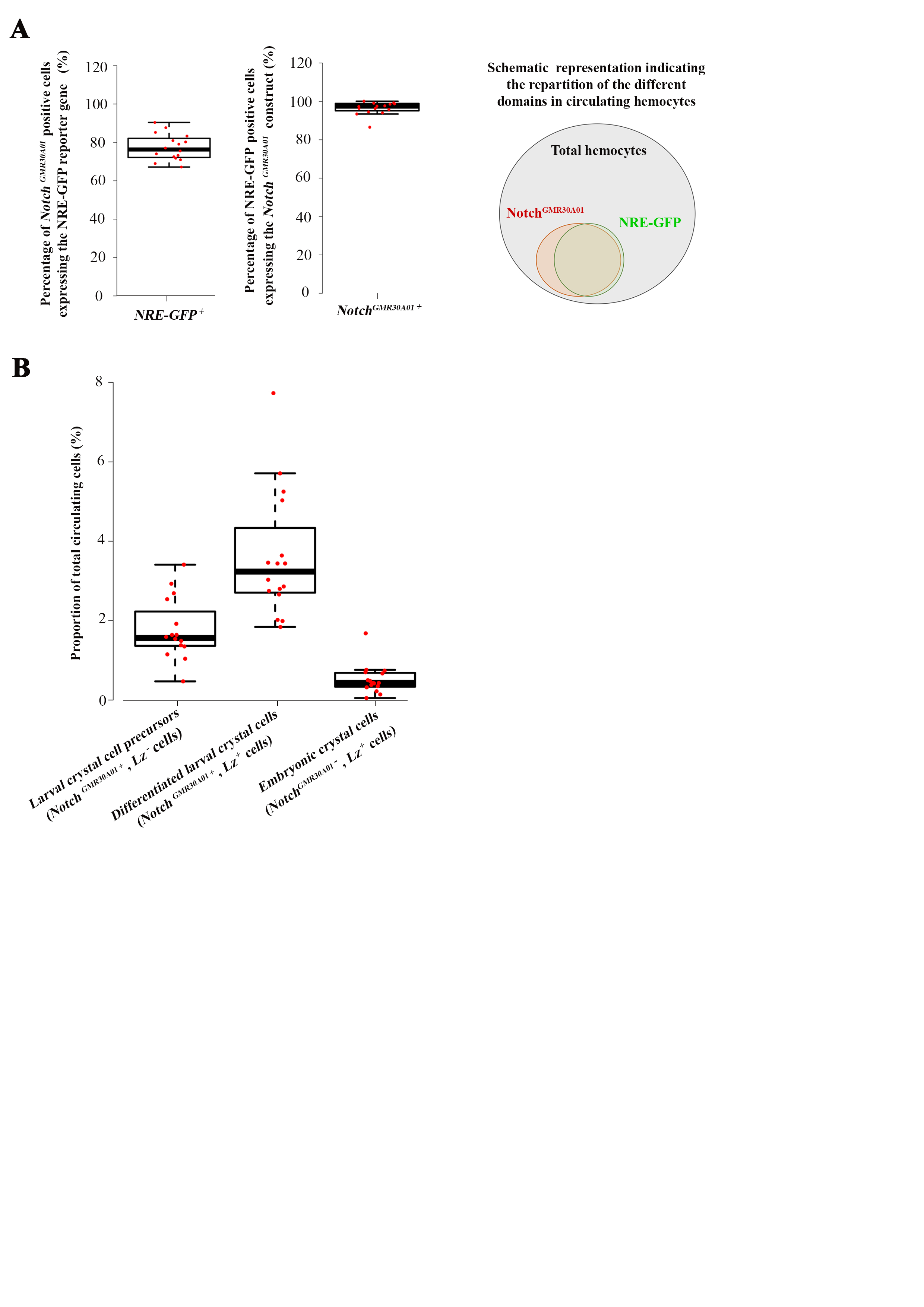

### Supplemental Figure 3

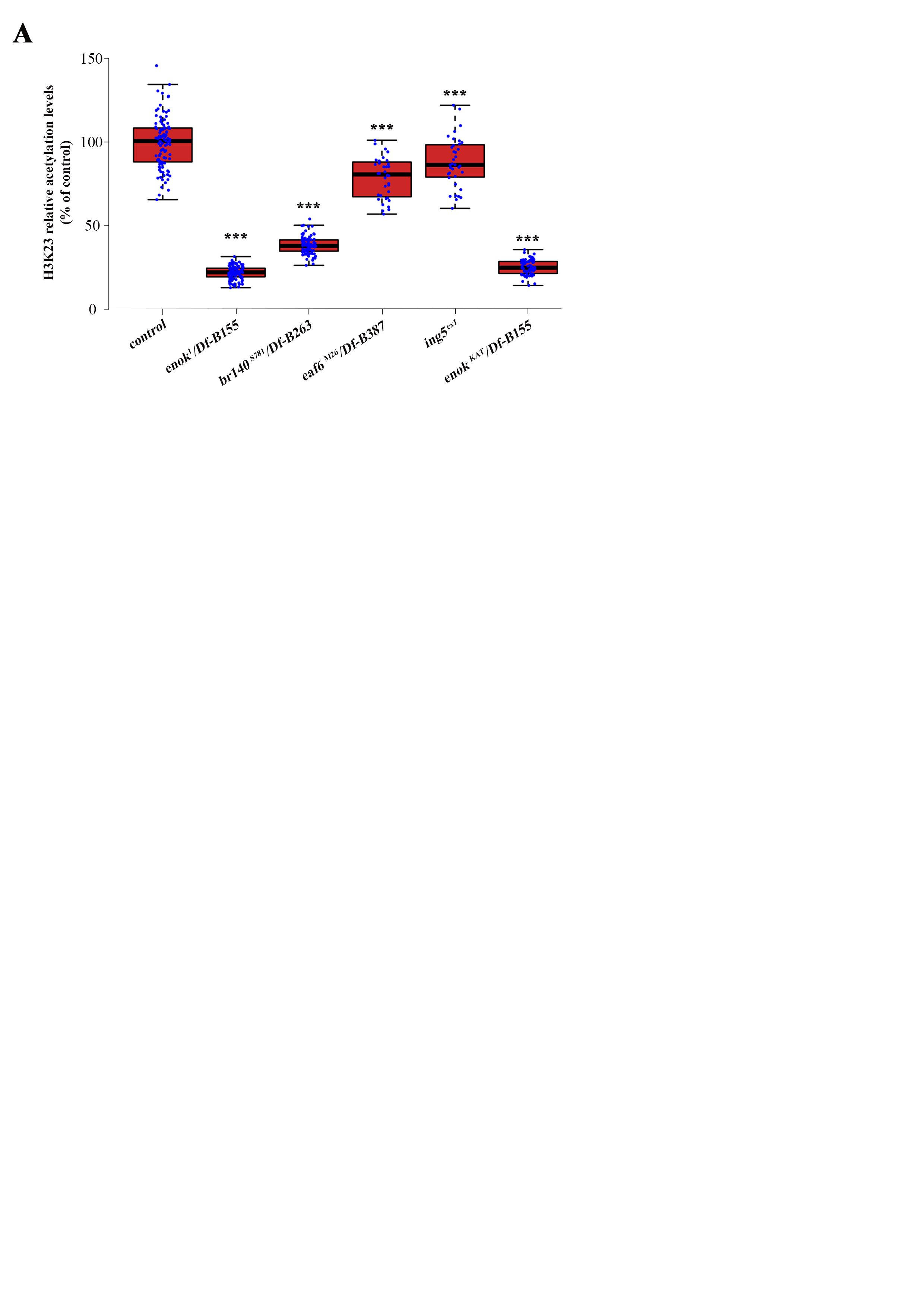

### Supplemental Figure 4

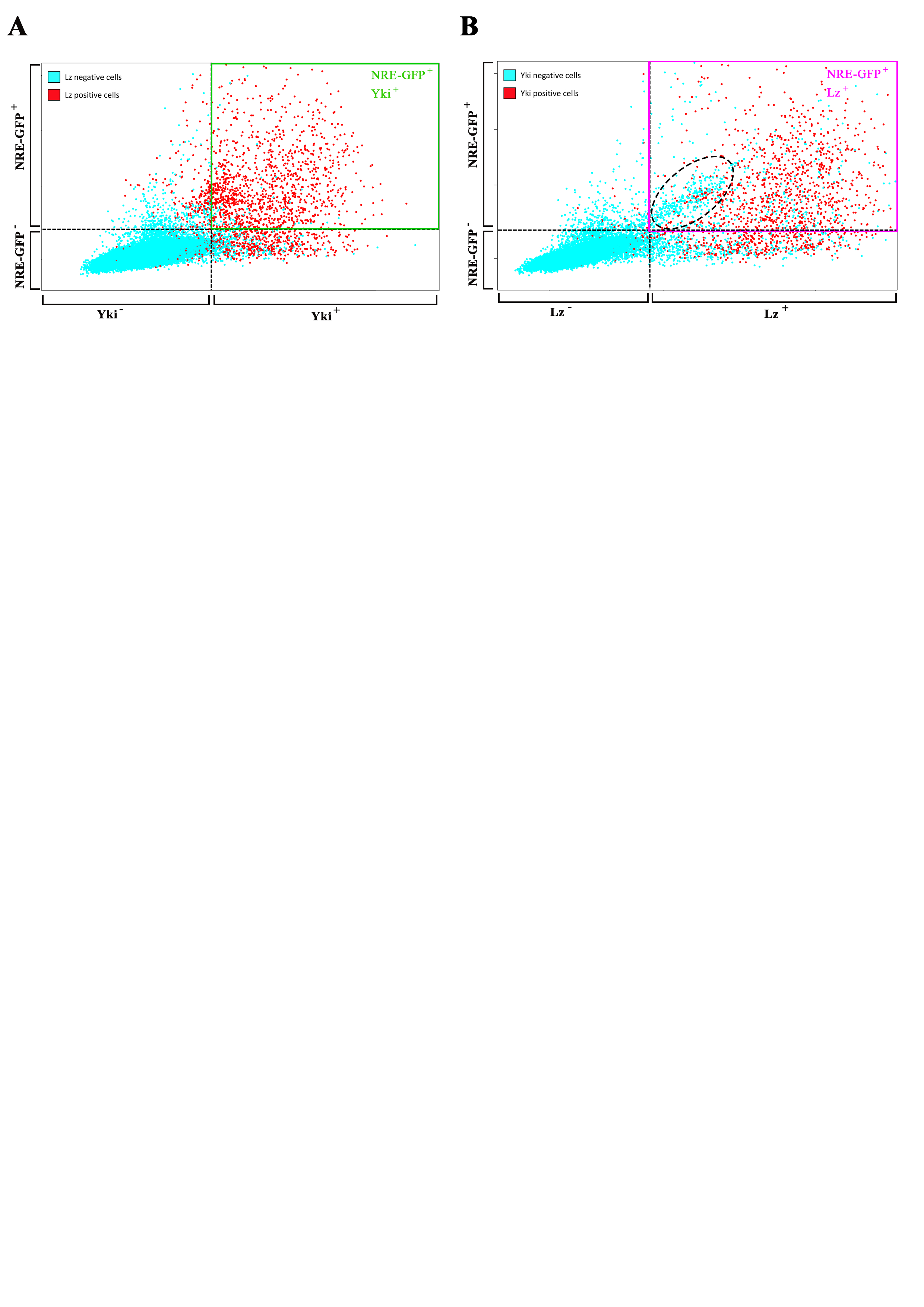

### Supplemental Figure 5

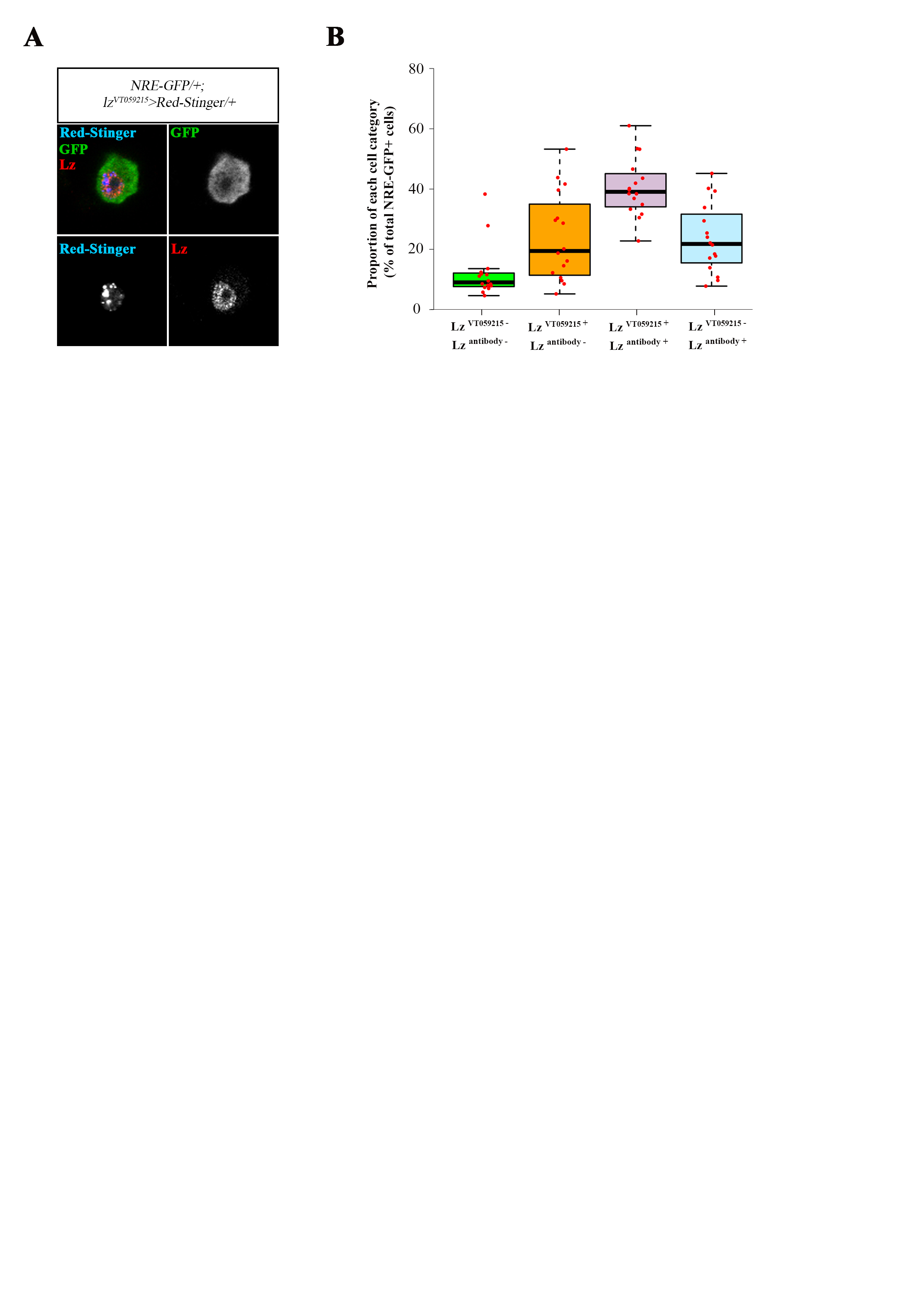
